# UBP12 and UBP13 deubiquitinases destabilize the CRY2 blue-light receptor to regulate growth

**DOI:** 10.1101/2021.04.29.441934

**Authors:** Louise Norén Lindbäck, Oliver Artz, Amanda Ackermann, Ullas V. Pedmale

## Abstract

All organisms undergo growth, which is precisely controlled by exogenous and endogenous signals. Unchecked growth often leads to neoplasia and other developmental defects, severely affecting an organism’s fitness. Light is a vital exogenous signal sensed by cryptochrome (CRY) blue light receptors to modulate growth and the circadian clock in plants and animals. Yet, how CRYs interpret light quantity to regulate growth in plants remains poorly understood. We show that UBP12 and UBP13 deubiquitinases physically interact with CRY2 in light. UBP12/13 negatively regulated CRY2 protein levels by promoting its ubiquitination and turnover to fine-tune growth. Unexpectedly, the destabilization of CRY2 by UBP12/13 is contrary to the general view that deubiquitinases stabilize proteins by preventing their degradation. Growth and development were explicitly affected in blue light when UBP12/13 was disrupted or overexpressed, indicating their role alongside CRY2. UBP12/13 also interacted and stabilized COP1, which is partially required for the turnover of CRY2. Despite decades of studies on deubiquitinases, the knowledge on how they are regulated is limited. Our study offers an insight into how exogenous signals and their receptors regulate deubiquitinase activity by protein-protein interaction. Altogether, our results provide a new module of cryptochromes and deubiquitinases in sensing and interpreting light cues to control growth at the most appropriate time.

## INTRODUCTION

Multicellular organisms undergo growth and development during their lifetime within the scope of their genetic and developmental constraints. However, the mechanisms by which growth and development are regulated have been elusive. Unchecked growth often leads to neoplasia in animals and loss of fitness in plants and other species. Therefore, understanding growth has the potential to prevent and cure cancer and provides an opportunity to improve crop productivity and biomass. Growth is either restricted or promoted at the most appropriate time based on many endogenous and exogenous signals. Light is one such important signal utilized by organisms to regulate their growth and physiology. In animals, light affects mood, behavior, metabolism, growth, and entrainment of the circadian clock (Fernandez et al. 2018; Bedrosian and Nelson 2017). Light is a source of energy in plants and an endogenous signal to module their growth and development, inform them about their geographical location on the planet, enable them to change their body plan to better adapt to their environment, and entrain their biological clocks (Chory 2010).

Apart from rods and cones that impart vision to the organism, there are specialized photoreceptors that perceive specific ultraviolet (UV), blue, red, and far-red wavelengths to monitor their quality and quantity to inform the organism’s environment’s state. UV-A/blue light-absorbing cryptochromes (CRY) are one such evolutionary conserved photoreceptors found in broad lineages including yeast, flies, plants, and animals (Pedmale et al. 2016; Sancar 2003). CRYs evolved from DNA photolyases that utilize UV-A/blue light as an energy source to repair damaged DNA. Present-day CRYs have retained the light absorption properties of photolyases but can no longer bind DNA directly. Two copies of CRYs are generally present in mammals and plants, namely CRY1 and CRY2, which are known to entrain the circadian clock, regulate metabolism, and control other crucial processes in them. Loss of animal CRYs is associated with tumorigenesis, diabetes, and neuronal disorders (Hirota et al. 2012; Lamia et al. 2011). The major roles of plant CRY1 and CRY2 are their regulation of light-dependent development termed photomorphogenesis, control of seedling stem (hypocotyl) growth, and photoperiodic flowering. CRYs are able to modulate these wide ranges of processes by governing gene expression through their interaction with signaling partners, often with bHLH transcription factors like BMAL1 (Brain and Muscle ARNT-Like 1) and CLOCK (Circadian Locomotor Output Cycles Kaput) in animals, whereas with PIFs (Phytochrome Interacting Factors) and CIBs (Cryptochrome-Interacting Basic helix-loop-helixes) in plants (Pedmale et al. 2016; Ma et al. 2016; Koike et al. 2012; Liu et al. 2008a). Central to CRY-mediated signaling in plants, animals, flies are their targeted ubiquitination and degradation by the 26S proteasome. Proteasomal degradation of CRYs is hypothesized to be necessary for their desensitization and reset their downstream signaling (Liu et al. 2016; Godinho et al. 2007).

The mechanisms underlying light control of growth, primarily through CRY photoreceptors, have been elusive in plants. In *Arabidopsis*, CRY1 is a nuclear and cytoplasmic localized protein, while CRY2 is nuclear and forms punctate nuclear speckles after blue light absorption (Yu et al. 2007). Photoactivated CRYs oligomerize as a tetramer, which is understood to be its physiologically active form (Ma et al. 2020; Palayam et al. 2021). Heterodimerization of CRY2 with BIC1 (Blue-light Inhibitor of Cryptochromes 1), interferes with its oligomerization to inactivate it (Wang et al. 2016). One of the roles of photoactive CRY2 is to dampen the activity of COP1-SPA (constitutive photomorphogenic 1-suppressor of PhyA-105) ubiquitin ligase, which serves as a substrate-specific adaptor of the Cullin 4-RING ubiquitin ligase (CRL4). CRL4^COP1-SPA^ promotes ubiquitination and subsequent degradation of transcription factors such as HY5, HYH, and LAF1 (Seo et al. 2003; Liu et al. 2011). Inactivation of CRL4^COP1-SPA^ by CRY2 leads to accumulation of these transcription factors and contributes to plant development and growth. CRY2 protein abundance is also regulated by light, accumulating in darkness or the vegetational shade, and rapidly turned over under prolonged and high blue light intensities (Pedmale et al. 2016; Yu et al. 2007).

Therefore, CRY2 protein levels and activity are tightly regulated to fine-tune hypocotyl growth and photomorphogenesis in blue light. In animals, F-box protein FBXL3 (F-box/LRR-repeat protein 3) functions as an E3 ubiquitin ligase to facilitate ubiquitination and degradation of mammalian CRY1 and CRY2 (Siepka et al. 2007; Busino et al. 2007; Godinho et al. 2007). Recently, it was shown CRL4^COP1-SPA^ and CRL3^LRBs^ are required for the degradation of CRY2 in *Arabidopsis* (Weidler et al. 2012; Liu et al. 2016; Lau et al. 2019; Ponnu et al. 2019; Chen et al. 2021). But the mechanism of action on how photoactive CRY2 is ubiquitinated by COP1 in light remains poorly understood, as they interact both in the dark and light and irrespective of CRY2’s phosphorylation status (Wang et al. 2001; Chen et al. 2021).

Protein ubiquitination is a key reversible post translational modification similar to phosphorylation and methylation. Polyubiquitination of proteins mostly serves as a signal for 26S proteasomal degradation, while monoubiquitination is often linked with non-degradation independent roles such as protein trafficking. E3 ubiquitin ligases ubiquitinate their target proteins with high specificity to either cause their degradation, alter their activity or to affect their localization. Ubiquitination can be reversed by deubiquitinating enzymes (DUB), to generally prevent the targeted protein from degradation. DUBs are evolutionary conserved group of proteases that counter the action of E3 ubiquitin ligases by trimming the ubiquitin chains and/or removing ubiquitin covalently bound to the proteins (Komander et al. 2009). In both plants and animals, DUBs comprise of five major gene families: ubiquitin-specific proteases (UBP/USP), ubiquitin-carboxyl terminal (UCH) proteases, the ovarian tumor proteases (OUT), the Machado-Joseph disease protein domain proteases (MJD) and the Jab1/MPN^+^/MOV34 (JAMMs) proteases (March and Farrona 2018; Komander et al. 2009; Lai et al. 2020). However, unlike E3 ligases, the role of DUBs in mediating key cellular process is slowly emerging, especially in plants. Also, the molecular functions and the substrates of the large number (∼64) of plant DUBs remain largely unidentified.

CRY2 protein levels and activity are critical to plant growth in light, especially during photomorphogenesis, which guides newly germinated seedlings to establish and ensure success as an organism. The detailed mechanisms that account for the precise control of CRY2 protein level by ubiquitination to control plant growth is not fully understood. Therefore, we took advantage of plants’ phenotypic plasticity to understand the molecular mechanisms on how CRYs regulate growth. The hypocotyl growth is particularly sensitive to light intensity, they respond appropriately by growing longer or shorter based on light conditions. In this study, we identify UBP12 and UBP13 deubiquitinase-mediated regulation of CRY2 degradation as a mechanism to regulate hypocotyl growth in light. Unexpectedly and differing from the current documented belief, the critical function of UBP12/13 DUBs in this process, however, turned out not to depend on its deubiquitination activity to stabilize and prevent CRY2 from degradation. But instead, these DUBs used their influence to decrease CRY2 protein level by ubiquitin-mediated degradation. Hypocotyl growth was disrupted in seedlings lacking UBP12 and UBP13 or when they were overexpressed, specifically in blue light. Our combined genetic and molecular data support a mechanistic model where photoactivated CRY2 interacts directly with UBP12/13 DUBs, and this complex recruits COP1 by direct contact. Furthermore, UBP12/13 mediated deubiquitination of COP1 leads to its stabilization, which then promotes ubiquitination and degradation of CRY2 in blue light. This mechanism of attenuation of CRY2 is unusual among the reported mechanisms but likely typifies the receptor’s mitigation in an everchanging light environment of the plant. Such regulation by CRY2-UBP12/13-COP1 axis is particularly essential to optimize hypocotyl growth during photomorphogenesis.

## RESULTS

### UBP13 through its MATH domain interacts with the CRY2 receptor

To identify novel regulators of CRY2 protein abundance in plants, we examined the CRY2 protein complex in *Arabidopsis* seedlings by affinity purification coupled with mass-spectrometry. We used the Flag-CRY2 expressing transgenic lines that complements the *cry2* mutant (Pedmale et al. 2016) under a subdued blue light condition to isolate the CRY2 protein complex. Liquid chromatography-tandem mass spectrometry (LC-MS) analysis identified previously known interacting partners of CRY2, including COP1, SPAs, CRY1, and BIC1 (Figure S1A) (Wang et al. 2001; Weidler et al. 2012; Wang et al. 2016). Among these were also ubiquitin-specific protease (UBP) 12 and 13, which are DUBs that remove ubiquitin modifications to stabilize their target proteins (Figure 1A-1B). In *Arabidopsis*, UBP12 and UBP13 share 91% amino acid sequence identity suggesting their biological function is likely redundant (Figure S1B) (Cui et al. 2013). The orthologs of UBP12 and UBP13 proteins can be found in other plant species, invertebrates and vertebrates (Figure S2A) suggesting evolutionary conservation. Previously, UBP12/UBP13 has been shown to have roles in immunity, flowering, JA signaling, and leaf development (Cui et al. 2013; Vanhaeren et al. 2020; Jeong et al. 2017; Lee et al. 2019).

**Figure 1.**
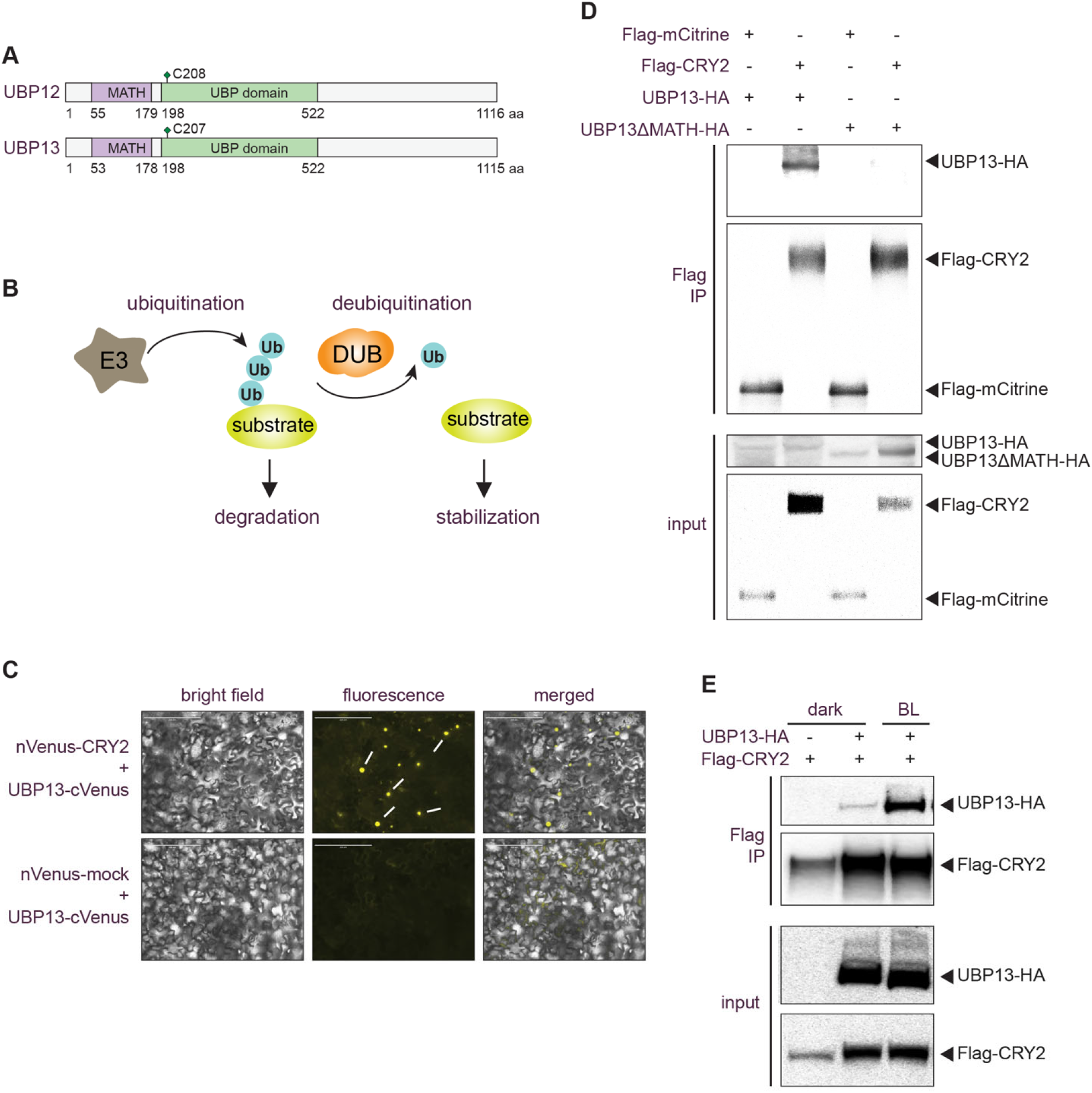
UBP13 physically interacts with CRY2 and their contact is enhanced in light. **(A)** Schematic representation of the *Arabidopsis* UBP12 and UBP13 deubiquitinase protein indicating the position of the conserved MATH and UBP domains containing the active cysteine catalytic residue. **(B)** Roles of E3 ubiquitin ligases (E3) and deubiquitinases (DUB) in controlling the fate of the target proteins by catalyzing their ubiquitination or deubiquitination. Ub indicates ubiquitin. **(C)** CRY2 and UBP13 physically interact in the nucleus as determined by bimolecular fluorescence complementation (BiFC) assay in *N. benthamiana*. The arrows indicate representative fluorescence reconstitution in the nucleus of the leaf epidermal cells. **(D)** *In planta* co-immunoprecipitation (co-IP) reveals that Flag-CRY2 interacts with UBP13-HA but not with UBP13 lacking the MATH domain (ΔMATH). Co-IP was performed using anti-Flag antibody and the immunoblot was performed using anti-HA and Flag antibodies. Flag-mCitrine serves as a negative control. **(E)** Co-immunoprecipitation (co-IP) analysis indicates that blue light enhances CRY2-UBP13 interaction in *Arabidopsis*. Co-IP was performed on protein lysates from *Arabidopsis* transgenic seedlings co-expressing UBP13-HA and Flag-CRY2. Seedlings were either blue light (30 µmol m^- 2^s^-1^) or mock treated in darkness. Co-IP was performed using anti-Flag antibody, and anti-HA and anti-Flag antibodies were used for the immunoblot. *Arabidopsis* seedlings expressing only Flag-CRY2 was used as a negative control.

First, we validated the CRY2-UBP13 interaction *in vivo* using bimolecular fluorescence complementation (BiFC). We found that CRY2 interacted with UBP13 in the nucleus of the epidermal cells (Figure 1C), since UBP12 and UBP13 are nucleocytoplasmic localized (Cui et al. 2013; Derkacheva et al. 2016) whereas CRY2 is a nuclear protein (Yu et al. 2007). *Arabidopsis* encodes approximately 64 deubiquitinases, which comprises of 27 UBP gene family members (Liu et al. 2008b). Among them, only UBP12 and UBP13 contain the meprin and TRAF (MATH) domain which aids in protein-protein interactions especially with diverse receptors (Figures S1B and S2B) (Liu et al. 2008b; Ye et al. 1999). To test the role of MATH domain in CRY2-UBP13 interactions, we immunoprecipitated Flag-CRY2 from plant protein extracts that expressed HA-epitope tagged UBP13 (UBP13-HA) or UBP13 without its MATH domain (UBP13ΔMATH-HA) (Figures 1A and S2B). Immunoblotting for HA revealed that CRY2 failed to co-immunoprecipitate with UBP13 lacking its MATH domain (Figure 1D), and we validated this was not due to inadvertent changes in subcellular localization and thus loss of interaction with CRY2 (Figure S2C).

We next addressed if light influenced the interaction between CRY2 and UBP13. We performed co-immunoprecipitation using transgenic *Arabidopsis* seedlings that expressed both Flag-CRY2 and UBP13-HA. 5-days old seedlings were dark-adapted for 24 hours, then exposed to blue light for 10 minutes (30 µmol m^-2^ s^-^1) or mock-exposed (dark). Protein extracts were immunoprecipitated with the anti-Flag antibody, and subsequent immunoblotting with anti-HA antibody revealed that the Flag-CRY2 interaction with UBP13-HA was greatly enhanced in blue light (Figure 1E). Together, these findings suggest that UBP12/13 are new signaling mediators in the CRY signaling pathway, and that UBP13 interacts with CRY2 in the nucleus through its MATH domain, and that blue light enhances their interaction.

### UBP12 and UBP13 controls hypocotyl growth and CRY2 protein level in blue light

When seedlings first emerge from the soil and encounter sunlight, they undergo photomorphogenesis, a series of development changes including cotyledon expansion, greening and hypocotyl growth inhibition (Chory 2010). CRY2 is known to mediate blue light-mediated hypocotyl growth inhibition under low fluence levels (<1 µmol m^-2^ s^-1^) (Lin et al. 1998). Therefore, we tested the effect of UBP12 and UBP13 on the blue light-mediated hypocotyl growth on seedlings grown for 4-days in blue light (1 µmol m^-2^ s^-^1) using single mutant alleles of UBP12 (*12-1* and *12-2w*) and UBP13 (*13-1* and *13-3*), along with *ubp12-2w ubp13-3* (*ubp12ubp13*) double mutant, *cry2*, and wild type (WT). Loss of either UBP12 or UBP13 in seedlings did not affect hypocotyl length (Figures 2A-2B) further signifying redundancy in their biological function (Figure S1B) and confirming a previous report (Cui et al. 2013). In contrast to the single mutants, *ubp12ubp13* double mutants developed a hypersensitive short hypocotyl phenotype in blue light, whereas *cry2* mutants had a blue light insensitive long hypocotyl phenotype (Figures 2A-2B).

**Figure 2.**
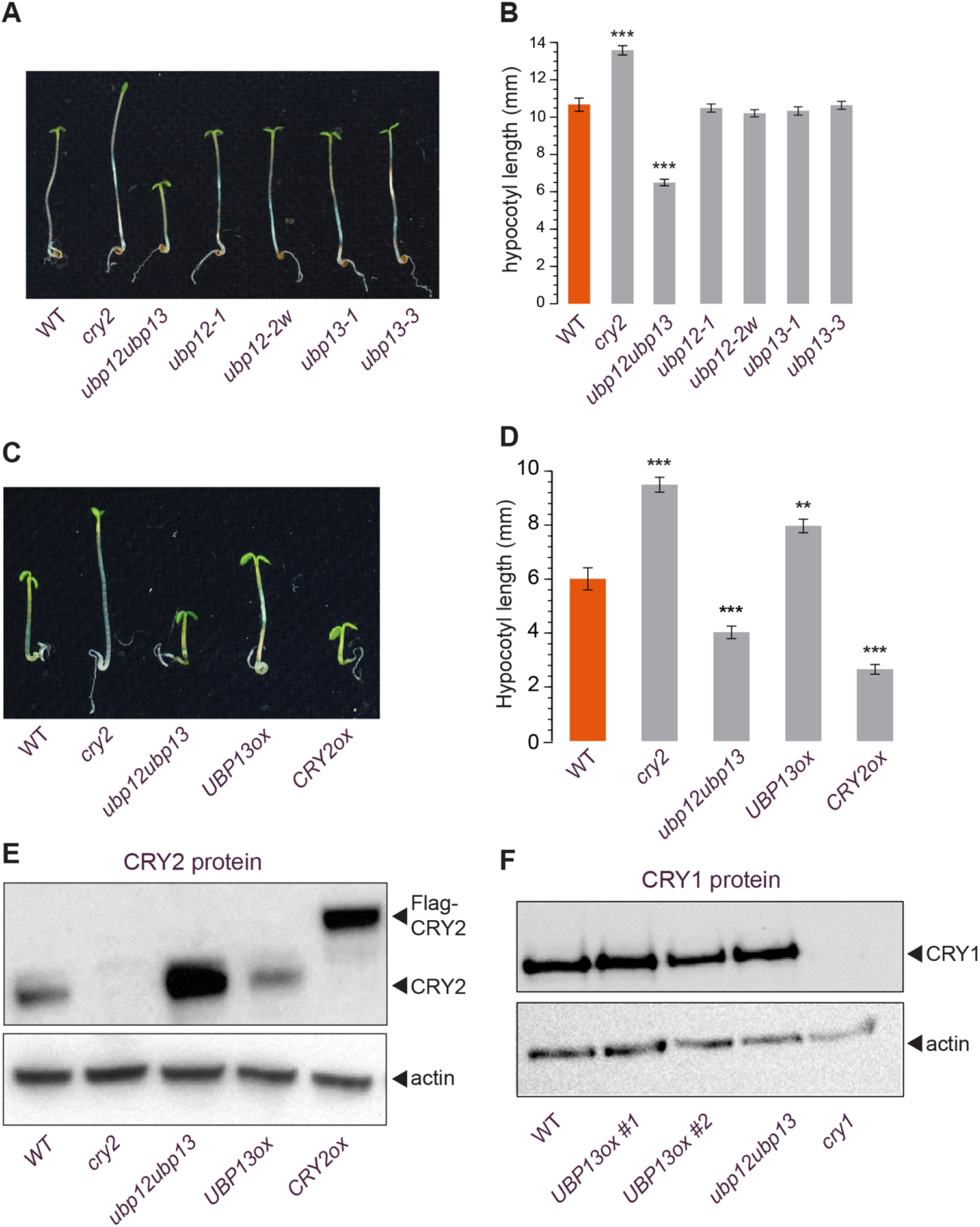
UBP12 and UBP13 negatively regulates CRY2 protein and controls growth in blue light. (**A-B**) Hypocotyl growth assay in blue light reveals that UBP12 and UBP13 regulate hypocotyl growth in blue light. Seedlings of the indicated genotypes were grown for 4 days in 1 µmol m^-2^s^-1^ constant blue light and their hypocotyl length was then measured. Single ubp12 and ubp13 mutants had no phenotype indicating redundancy, but *ubp12ubp13* double mutant showed significantly shorter hypocotyl phenotype compared to the WT. *cry2* mutant displayed the expected blue light insensitive long hypocotyl phenotype. ***, *p* < 0.001; Student’s t-test. Error bars represent standard error. (**C-D**) Hypocotyl growth assay indicates that UBP13 overexpressor (*UBP13ox*) had a blue light insensitive hypocotyl phenotype similar to *cry2* mutant. Whereas, the *ubp12ubp13* and *CRY2ox* seedlings had a short hypersensitive hypocotyl in blue light. The hypocotyl length of the indicated genotypes is indicated after growing them in blue light for 4 days. Student’s t-test, ***, *p* < 0.001, **, p < 0.01. **(E)** Immunoblot analysis reveals that UBP12 and UBP13 negatively regulates CRY2 protein in blue light. In *ubp12ubp13* seedlings, increased CRY2 protein accumulation is seen when compared with WT and CRY2 overexpressor tagged with Flash epitope tag (*CRY2ox*). Overexpression of UBP13 leads to lower CRY2 levels. Total protein lysates from 4-day-old seedlings grown in blue light (1 µmol m^-2^ s^-1^) were probed with anti-CRY2 antibody. Actin serves as a loading control. **(F)** UBP12 and UBP13 does not regulate CRY1 protein levels in blue light. Total protein lysates from 4-day-old seedlings grown in blue light (1 µmol m^-2^ s^-1^) were used and probed with anti-CRY1 antibody. Actin serves as a loading control.

These results indicate that UBP12 and UBP13 modulates hypocotyl growth in blue light. CRY2 protein levels regulate hypocotyl growth in blue light, their reduced levels result in a long hypocotyl phenotype, and higher levels lead to a shorter hypocotyl (Figure 2C-2D) (Lin et al. 1998; Pedmale et al. 2016). As UBP12 and UBP13 are deubiquitinases, we next investigated their role in regulating CRY2 protein levels and the hypocotyl growth in blue light. We measured hypocotyl growth of *ubp12ubp13* and *cry2* mutants, and CRY2 (*CRY2ox)* or UBP13 *(UBP13ox)* overexpressing seedlings grown for 4 days in blue light (1 µmol m^-2^ s^-1^). *ubp12ubp13* double mutants developed short hypocotyls, mimicking *CRY2ox* (Figures 2C-2D), whereas *UBP13ox* overexpression led to a long hypocotyl phenotype, comparable to *cry2* mutants only under blue light (Figures 2C-2D and S3A-S3B). This blue light specific phenotype of *UBP13ox* and *ubp12ubp13* was not observed in red or white light (Figures S3C-S3D). To assess if UBP12 and UBP13 regulate CRY2 levels in blue light, we performed immunoblot analysis on protein extracts obtained from these genotypes using an anti-CRY2 antibody. Notably, CRY2 protein levels were much higher in *ubp12ubp13* seedlings than WT and *CRY2ox* seedlings, whereas the *UBP13ox* had a reduced amount of CRY2 (Figure 2E) specific in blue light as we did observe the same in other light conditions (Figures 3F-3G). Though we did not detect a dramatic prominent decrease in CRY2 in *UBP13ox* when compared to the *ubp12ubp13* seedlings, the lack of hypocotyl growth inhibition was evident in multiple independent transgenic lines (Figures S3A-S3B, S3E). The effect of UBP12 and UBP13 was also specific to CRY2, as we did not see changes in CRY1 protein levels as determined by immunoblot assay using an anti-CRY1 antibody (Figure 2F). The unexpected result that UBP12 and UBP13 negatively regulate CRY2 protein levels is opposite of the expectation of a deubiquitinase. As deubiquitinases are generally known to deubiquitinate and prevent the degradation of their target proteins (Komander et al. 2009; March and Farrona 2018), one would hypothesize and expect that the loss of UBP12 and UBP13 will lead to lower CRY2 protein level resulting in a longer hypocotyl. Similarly, overexpression of UBP12 or UBP13 will lead to increased CRY2 protein stabilization resulting in a shorter hypocotyl as seen in *CRY2ox* seedlings. Here, our results suggest that UBP12 and UBP13 have a contrary effect, destabilizing CRY2 to specifically regulate CRY2 protein amount and CRY2-dependent hypocotyl inhibition in blue light.

**Figure 3.**
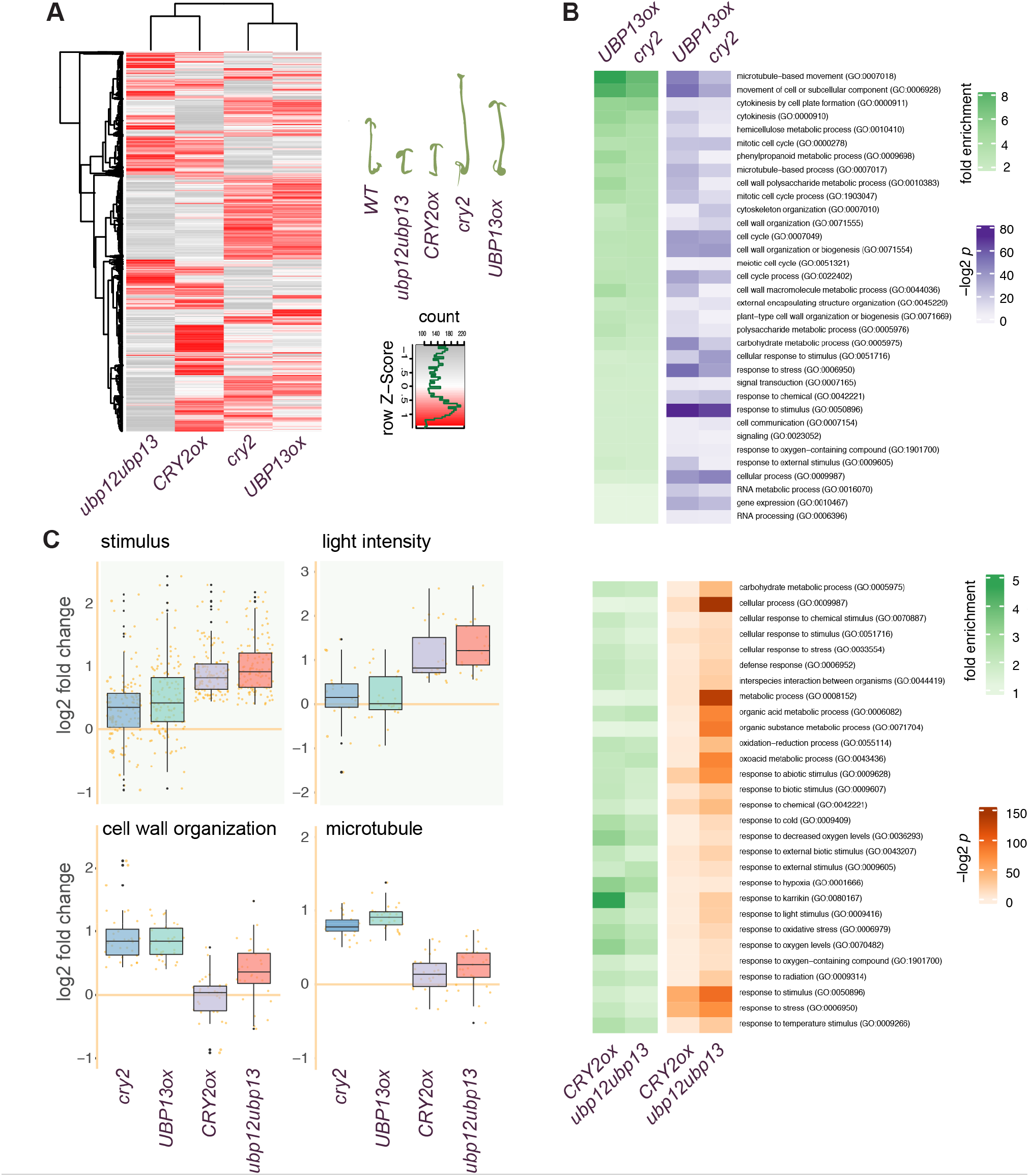
Genotypes with similar growth phenotype have similar gene expression patterns. **(A)** Unbiased hierarchical clustering of the differentially expressed genes compared to the WT. This illustrates a shared gene expression pattern between *ubp12ubp13* and *CRY2ox*, which has a short hypocotyl phenotype. Similarly, between *cry2* and *UBP13ox*, which has a long hypocotyl. RNA-seq analysis was performed on mRNA isolated from 4-day-old seedlings of the indicated genotypes grown in 1 µmol m^-2^ s^-1^ blue light. **(B)** Top and common (ranked by fold enrichment) of significantly enriched GO terms derived from genes that are upregulated in *ubp12ubp13* and *CRY2ox*, which has short hypocotyl phenotype in blue light, and between *cry2* and *UBP13ox*, which has a long hypocotyl phenotype **(C)** Representative expression patters of the indicated gene families in the specified genotypes. Center lines are the medians; box limits indicate the 25^th^ and 75^th^ percentiles, whiskers extend 1.5 times the interquartile range from the 25^th^ and 75^th^ percentiles; outliers are represented by black dots.

### Genotypes with similar hypocotyl growth phenotypes have similar gene expression patterns

Next, to corroborate whether the genotypes exhibiting similar long hypocotyl phenotypes (*UBP13ox* and *cry2)* and short hypocotyl phenotypes (*ubp12ubp13* and *CRY2ox*) also caused similar gene expression changes, we analyzed their transcriptome in blue light. We performed mRNA-seq analysis on WT, *cry2, ubp12ubp13, CRY2ox*, and *UBP13ox* seedlings grown in blue light (1 µmol m^-2^ s^-1^) for 4 days using biological replicates which correlated with each other, with Pearson *R* ranging between 0.98-0.99 (Figure S4A). We identified significantly differentially expressed genes (DEG; false discovery rate, FDR *<0*.*05*) in comparison to the WT (table S1). Unbiased hierarchical clustering of these DEG identified two nodes, one belonging to *ubp12ubp13* and *CRY2ox;* and the other for *cry2* and *UBP13ox* (Figure 3A). Gene ontology (GO) analysis further supported that *cry2* and *UBP13ox* have comparable gene expression changes, and likewise between *ubp12ubp13* and *CRY2ox* (Figure 3B). We found genes responding to light stimulus and intensity were over-represented in *CRY2ox* and *ubp12ubp13* mutants (Figure 3C), indicating active light-dependent signaling and gene expression leading to hypocotyl inhibition (Pedmale et al. 2016; Huang et al. 2019). In contrast, genes involved in cell-wall organization or in microtubule assembly, characteristic of a growing or uninhibited hypocotyl (Pedmale et al. 2016; Sasidharan and Pierik 2010), were upregulated in *cry2* mutants and *UBP13ox* (Figure 3C). CRYs in animals and plants, mostly modulate gene expression through its interaction with bHLH family of transcription factors, which bind to the E/G-box elements present in the promoter sequence of the genes (Koike et al. 2012; Pedmale et al. 2016). Therefore, we reasoned that increased CRY2 activity will result in expression of genes with overrepresentation of G-box cis-elements in their promoters. Indeed, among the promoter sequence of the common genes upregulated in *CRY2ox* and *ubp121ubp13*, we identified G-box [CACGTG] cis-motif but not in *cry2* and *UBP13ox* genotypes (Figure S4B). These observations indicate that physiological and phenotypic changes due to changes in gene expression are sensitive to CRY2 protein levels being regulated in blue light. Furthermore, the findings further reinforce that UBP13 is involved in negatively regulating CRY2 and its function to control hypocotyl growth in blue light.

### UBP12 and UBP13 is required for the ubiquitination and degradation of CRY2

To gain further insight into the role of UBP12 and UBP13 in regulating CRY2 protein levels, we generated a triple mutant (*cry2ubp12ubp13*) by genetically crossing *ubp12ubp13 and cry2*. The hypocotyl length of the *cry2ubp12ubp13* was similar to *cry2* single mutant (Figure 4A), indicating that *CRY2* is epistatic to *UBP12* and *UBP13*. This genetic evidence further supports that CRY2 and UBP12 and UBP13 function in the same pathway to regulate hypocotyl growth in blue light. As loss of UBP12 and UBP13 (*ubp12ubp13* mutant) elevates CRY2 protein (Figure 2E) and leads to hypersensitive short hypocotyl in blue light (Figure 2C-2D), we examined whether CRY2 overexpression in *ubp12ubp13* will further reduce its hypocotyl length. Transgenic seedlings overexpressing CRY2 in *ubp12ubp13* mutant exhibited enhanced hypocotyl inhibition compared to *CRY2ox* and *ubp12ubp13* seedlings (Figure 4A), which is indicative of increased CRY2 protein abundance and stabilization.

**Figure 4.**
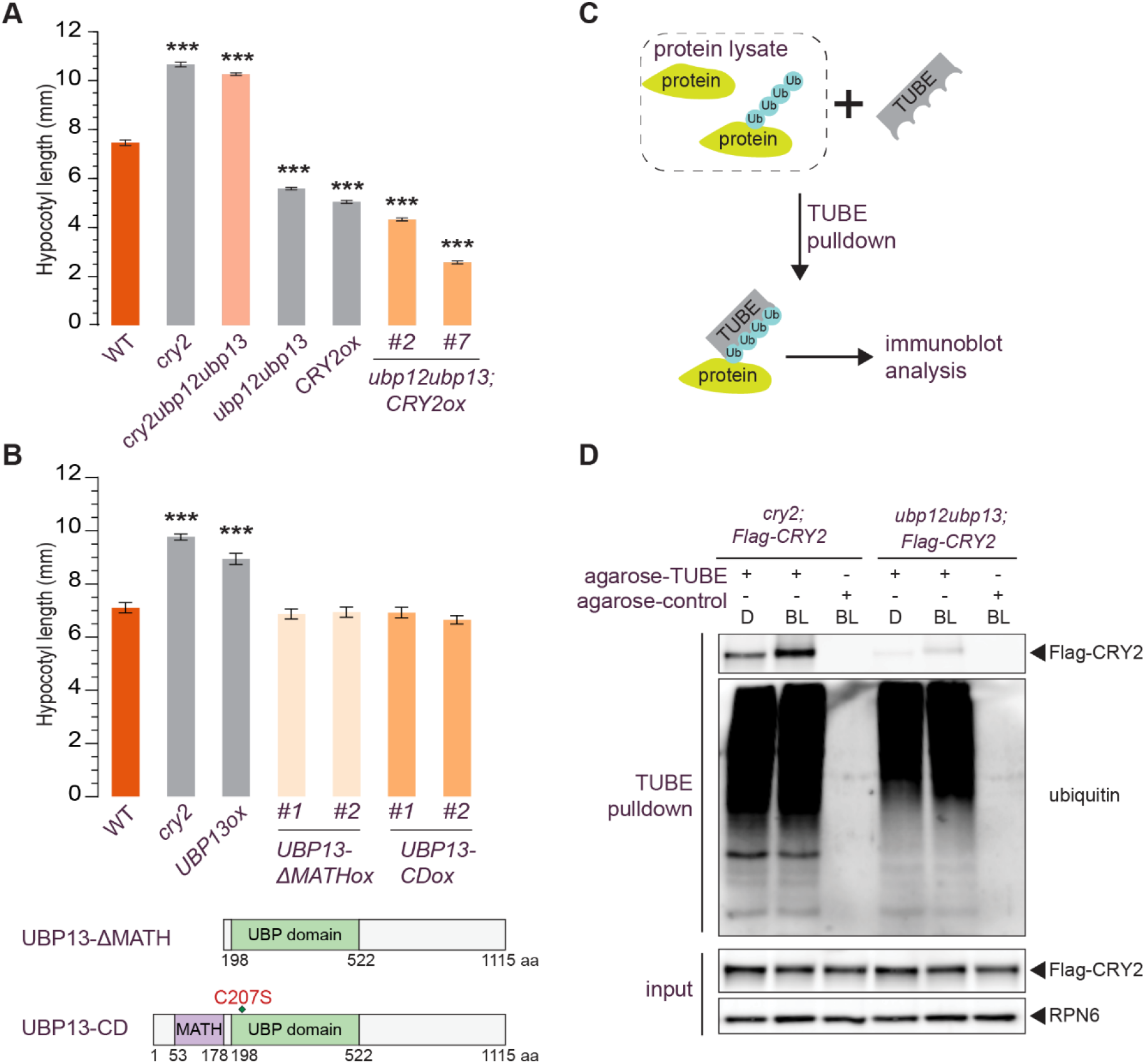
UBP12 and UBP13 is required for the ubiquitination and degradation of CRY2 in blue light. **(A)** Hypocotyl length analysis in blue light of the triple *cry2ubp12ubp13* mutant implies that CRY2 is epistatic to UBP12 and UBP13. Also, it signifies that CRY2 and UBP12/13 function in the same pathway in blue light. Overexpression of CRY2 in *ubp12ubp13* mutant seedlings causes much shorter hypocotyl than CRY2 overexpressor indicating that UBP12/13 regulate CRY2 protein. Hypocotyl length and phenotypes of the indicated genotypes after growing them in blue light (1 µmol m^-2^s^-1^) for 4 days is represented. **(B)** Deletion of the MATH domain or mutating the catalytic residue of the UBP13 had no obvious phenotypes. This indicates that the MATH domain and the catalytic activity of UBP13 is required to regulate CRY2 mediated hypocotyl growth in blue light. Hypocotyl length of seedlings of the indicated genotypes are shown after growing them for 4 days in blue light. *UBP13-ΔMATHox* transgenic line indicates deletion of the MATH domain, *UBP13-CDox* lines indicate mutation of the UBP13 with the catalytic active residue (C207S) resulting in a catalytically dead (CD) protein. Error bars represent standard error of the mean. Student’s t-test, ***, *p* < 0.001, **, p < 0.01. **(C)** Illustration of Tandem Ubiquitin Binding Entities (TUBE) pulldown method to enrich for ubiquitinated proteins. TUBE have a high affinity for polyubiquitylated proteins and enables efficient isolation and identification of ubiquitylated proteins from cell lysates. **(D)** UBP12 and UBP13 controls CRY2 ubiquitination in blue light. 4-day-old etiolated seedlings were either kept in darkness or treated with 30 µmol m^-2^s^-1^ blue light for 30 min before sample collection. Total ubiquitylated proteins were immunoprecipitated using agarose-TUBE from the *cry2;UBQ10pro::Flag-CRY2* and *ubp12ubp13;UBQ10pro::Flag-CRY2* seedlings in the dark and blue light treated samples. Agarose-control beads without TUBEs served as a negative control. The samples were analyzed by immunoblotting with anti-Flag antibody to detect CRY2, anti-ubiquitin antibody (P4D1) for detection of ubiquitylated proteins and RPN6 served as a loading control.

When we overexpressed UBP13-ΔMATH in *Arabidopsis* transgenic lines, there was no hypocotyl length phenotype, reinforcing that the MATH domain of UBP13 that mediates the contact with CRY2 is required for the hyposensitive phenotype seen in *UBP13ox* (Figures 4B and S5A). We also tested if the enzymatic activity of UBP13, that catalyzes the hydrolysis of the bonds between the ubiquitin and its substrate, affects CRY2 mediated hypocotyl growth. We generated transgenic lines that overexpress catalytically-dead UBP13 (*UBP13-CDox)*, in which the cysteine at position 207 in the conserved catalytic box (Cui et al. 2013) was changed to serine (Figure S2B).

Independent transgenic *UBP13-CDox* expressing seedlings were identical to the WT seedlings signifying that the deubiquitinase activity of UBP13 is required for CRY2 mediated hypocotyl growth (Figures 4B and S5A). Collectively, these results indicate that the physical interaction of UBP13 with CRY2 and the enzymatic activity of UBP13 are required to regulate CRY2 protein abundance and subsequently hypocotyl growth in blue light.

### CRY2 stability and ubiquitination is regulated by UBP12/13

Since our results indicate that UBP12 and UBP13 deubiquitinases and CRY2 function in the same pathway, and UBP12/13 negatively regulate CRY2 levels, we tested whether UBP12/13 controls the ubiquitination of CRY2. Dark-grown seedlings expressing Flag-CRY2 complementing the *cry2* mutant (*cry2; Flag-CRY2)*, or in the *ubp12ubp13* mutant background (*ubp12ubp13; Flag-CRY2)*, were exposed to blue light (30 µmol m^-2^ s^-1^) for 30 minutes or kept in the dark before total proteins were extracted. We then captured ubiquitinated proteins from these protein extracts using the Tandem Ubiquitin Binding Entities (TUBE) method (Figure 4C) (Hjerpe et al. 2009) followed by immunoblotting with anti-Flag antibody to detect CRY2 (Figure 4D). Immunoblotting with anti-ubiquitin antibody confirmed that ubiquitinated proteins were purified from both the dark and blue light treated seedlings (Figure 4D). Consistent with the previous reports that blue light stimulates ubiquitination of CRY2 (Liu et al. 2016), we observed enrichment of ubiquitinated Flag-CRY2 in blue light treated *cry2; Flag-CRY2* seedlings. However, we saw little ubiquitination of Flag-CRY2 in *ubp12ubp13* mutant seedlings in blue light (Figure 4D). Our findings here indeed suggest that UBP12 and UBP13 controls CRY2 ubiquitination and subsequent protein levels in blue light.

### UBP13 interacts with COP1 E3 ubiquitin ligase and increases its stability to mediate the degradation of CRY2

Our data so far indicates that the UBP12/13-CRY2 complex likely regulates the activity of another protein, which in turn destabilizes CRY2 under blue light. We reasoned that one such candidate is the COP1 E3 ubiquitin ligase which is part of the CRL4^COP1-SPA^ complex. Previous studies have shown that COP1 is partially required for the turnover of CRY2 (Figure S5B) (Wang et al. 2001; Lau et al. 2019; Ponnu et al. 2019; Weidler et al. 2012; Wang et al. 2015; Chen et al. 2021). Furthermore, COP1 and SPAs interact with CRY2, and we also identified them in the same protein complex along with CRY2, UBP12, and UBP13 (Figure S1A). However, it is not known how the partial turnover of CRY2 is mediated by COP1-SPA. Like many other E3 ligases, COP1 undergoes auto-ubiquitination to regulate its levels (Seo et al. 2003). Therefore, we tested if UBP12/13 affects COP1 protein levels in blue light. We indeed found that UBP13-HA with or lacking its MATH domain interacted with COP1-Flag (Figure 5A), preferentially in the nucleus as determined by BiFC (Figure S5C). This suggests that the MATH domain, which is crucial for UBP13-CRY2 interaction, is not required for the contact between UBP13 and COP1. We next determined COP1 protein levels in *ubp12ubp13, UBP13ox*, and COP1 overexpressor tagged with Flag (*COP1ox*) seedlings grown in blue light. Immunoblotting analysis with anti-COP1 antibody showed that COP1 was present in much higher abundance in *UBP13ox*, and undetectable in the *ubp12ubp13* mutant (Figure 5B). This indicates that UBP12/13 stabilizes COP1 protein in blue light. We found that COP1 overexpressing (*COP1ox*) seedlings exhibited blue-light insensitive long hypocotyl phenotype similar to *cry2 and UBP13ox* seedlings, whereas the *cop1* mutant had a shorter hypocotyl (Figure 5C-5D). We also examined the hypocotyl growth of transgenic lines overexpressing COP1 in *ubp12ubp13* mutant in blue light. The long hypocotyl phenotype of *COP1ox* was not observed when UBP12/13 was absent in *ubp12ubp13* mutant (Figure 5E), indicating that UBP12 and UBP13 is required for the stability and activity of COP1. Taken together, these results suggest that UBP13 physically interacts with COP1 to stabilize it, possibly by deubiquitination, and imparts support that COP1 is required for the turnover of CRY2. Additionally, the results provide a mechanistic framework on how light perceived by CRY2, leads to its ubiquitination and turnover mediated by UBP12/13 stabilization of COP1.

**Figure 5.**
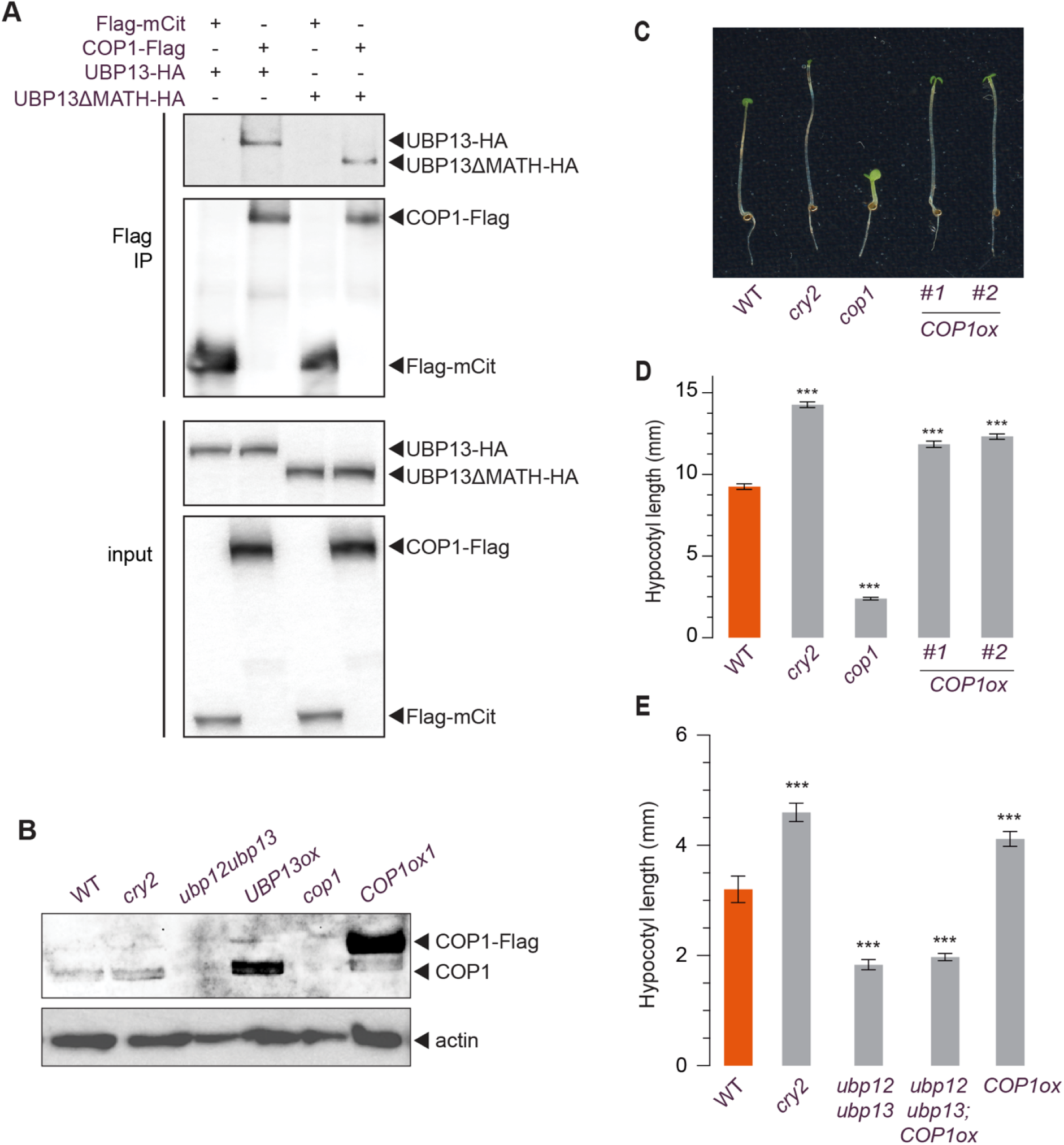
UBP13 interacts with COP1 and increases its stability to mediate degradation of CRY2 to regulate growth. **(A)** *In planta* co-immunoprecipitation (co-IP) reveals that COP1 and UBP13 physically interact, and the MATH domain of UBP13 that mediates UBP13-CRY2 interaction is not required for COP1-CRY2 contact. The UBP13-HA or UBP13ΔMATH-HA constructs were co-infiltrated with COP1-Flag into *N. benthamiana* leaves and immunoprecipitated with anti-Flag antibody. Immunoblot analysis was performed with anti-HA antibody to detect UBP13-HA. Flag-mCitrine was used as a negative control. **(B)** UBP12 and UBP13 positively regulates COP1 protein. In *ubp12ubp13* seedlings, COP1 protein is diminished like in the *cop1* mutant. Overexpression of UBP13 leads to higher COP1 levels compared to the WT. Total protein lysates from 4-day-old seedlings grown in blue light (1 µmol m^-2^ s^-1^) was used for this immunoblot and probed with anti-COP1 antibody. Actin serves as a loading control. A representative blot from at least three individual experiments is shown. (**C-E**) Seedlings of the indicated genotypes were grown for 4 days in 1 μmol m^-2^ s^-1^ constant blue light and their hypocotyl length was measured. Error bar indicates the standard error of the mean. Student’s t-test: ***, *p* < 0.001.

## DISCUSSION

Understanding growth and development not only answers a fundamental question about biological processes, but has broader applications in prevention and treating of cancer and other diseases and in increasing crop yield. However, the mechanisms that control an organism’s growth and size, particularly mediated by external stimuli, are not well understood. Light is a crucial external signal utilized by plants and animals for their normal function, such as synchronizing their circadian clocks, adjusting metabolism, and modulating their growth and development. However, we lack a complete comprehension of how organisms use light, especially perceived by CRYs, to control these processes. To answer this question, we took advantage of the plant’s phenotypic plasticity to study the mechanisms underlying growth. Due to the plants’ immobile nature, they are sensitive to their environment and readily modify their growth, create new or alter their body parts to better adapt to their surroundings. During seedling emergence from the soil, in a developmental process called photomorphogenesis, CRYs control seedling stem growth relative to the amount of blue light perceived by them. This control of hypocotyl growth depends on the amount of blue light; a higher amount of light results in degradation of CRY2, leading to attenuation of its signaling, and a lower light amount results in accumulation of CRY2 (Pedmale et al. 2016; Yu et al. 2007; Chen et al. 2021). CRY2 levels thus finetune growth in response to light levels; and so, it is essential to understand mechanisms that control CRY2 turnover.

Our study established a critical role of UBP12 and UBP13 DUBs in the CRY2 blue light signaling pathway to modulate seedling growth. UBP12/13 interacts with CRY2 and promotes its ubiquitination and turnover in blue light. DUBs generally oppose the action of E3 ubiquitin ligases, but UBP12/13 did not counteract the blue light-induced ubiquitination of CRY2, to stabilize and prevent its degradation. We further show that UBP12/13 controls the ubiquitination and degradation of CRY2 by direct interaction and stabilization of COP1. Consistent with this model, specifically in blue light, we show that the double *ubp12ubp13* and *cop1* mutants caused a short hypocotyl phenotype, similar to CRY2 overexpressing plants indicating increased CRY2 protein levels. UBP13 and COP1 overexpressing plants exhibited long hypocotyl phenotype, phenocopying *cry2* mutant indicating lower levels or loss of CRY2 protein. The turnover of CRY2 was accelerated in UBP13 overexpressing plants but was substantially stabilized and accumulated in *ubp12ubp13* mutant. Also, COP1 was unstable and barely detectable in *ubp12ubp13* double mutant, whereas it was stabilized when UBP13 was overexpressed. Together, our results reveal a key role for UBP12 and UBP13 in regulating hypocotyl growth and demonstrate a mechanism by which UBP12/13 regulates CRY2 abundance in blue light. This CRY2 abundance affects the sensitivity of the hypocotyl to blue light and thus modifies its growth.

UBP12/13 are functional deubiquitinases belonging to an ancient lineage that comprises of USP7/HAUSP (Ubiquitin Specific Protease 7/Herpesvirus-Associated Ubiquitin-Specific Protease) in metazoans and *Drosophila*, and the MATH-33 (MATH domain containing protein 33) in *Caenorhabditis elegans* (Figure S2A) (Heimbucher and Hunter 2015; Cui et al. 2013; Tian et al. 2012). *Arabidopsis* and human genome codes for 64 and 90 DUBs, in contrast, they have a large number of E3 ubiquitin ligases, 1400 in *Arabidopsis* and 700 in humans (Vierstra 2009; George et al. 2018; Lai et al. 2020; March and Farrona 2018). The lower number of DUBs suggests that each DUB can multitask and have a variety of substrates. This is indeed reflected from other studies in plants and animals, where UBP12/13 and USP7 regulate and are associated with multiple protein substrates functioning in many diverse signaling pathways. In plants, UBP12/13 constitutes the Polycomb repressive complex (PRC1) that regulates transcription (Lecona et al. 2015; Derkacheva et al. 2016). UBP12/13 modulates the jasmonate hormone pathway by binding to MYC2 transcription factor (Park et al. 2019); and associate with RGRF1/2 receptors during root development (An et al. 2018). DA1, DAR1 and DAR2 peptidases are deubiquitinated by UBP12/13 to modulate plant development (Vanhaeren et al. 2020). UBP12/13 interacts with ZEITLUPE (ZTL), a photoreceptor with ubiquitin ligase activity in the circadian clock (Lee et al. 2019). In animals, USP7 interacts with the tumor suppressor p53 and its ubiquitin ligase, MDM2. EBNA1 protein of Epstein-Barr Virus, commonly associated with human cancer, interacts with USP7 (Saridakis et al. 2005; Li et al. 2004). Furthermore, USP7 interacted with mammalian CRY1 and CRY2 alike to our study, but it deubiquitinated them, in contrast to our study.

Our unexpected finding that UBP12/13 negatively regulating CRY2 protein underscore a new and a previously undocumented role for a deubiquitinase. As DUBs are generally known to positively regulate proteins, UBP12/13 negatively regulated CRY2 by facilitating its ubiquitination and degradation, mirroring an E3 ubiquitin ligase’s function. This is in stark contrast to all the studies undertaken till date on UBP12/13, and its mammalian ortholog USP7. In previous studies and consistent with their function as a deubiquitinating protein, ectopic overexpression of USP7 and UBP12/13, stabilized their interacting partners. Conversely, their loss-of-function led to destabilization of their target proteins. For example, in mammals, p53, CRY1/2, MDM2 were stabilized when USP7 was overexpressed, and were destabilized and degraded when USP7 was ablated (Li et al. 2004, 2002; Papp et al. 2015). Similarly, *Arabidopsis* UBP12/13 positively controlled the stability of MYC2, ZTL, RGRF1/2, DA1, DAR1 and DAR2 proteins (Vanhaeren et al. 2020; An et al. 2018; Lee et al. 2019; Jeong et al. 2017).

We found that light enhanced CRY2 binding to the amino-terminal region of UBP13, containing the MATH domain to provide specificity. In addition, UBP13 interacted with the COP1 ubiquitin ligase at a region distinct from its MATH domain. DUBs are known to bind to ubiquitin ligases; for example, USP7 binds to MDM2 and RNF2/RING2, whereas UBP12/13 has been shown to interact with ZTL (Maertens et al. 2010; Li et al. 2002; Cummins et al. 2004). Interestingly, USP7 deubiquitinated MDM2, and its target p53, likewise, our data indicates that UBP12/13 can interact with CRY2 and its ubiquitinating ligase, COP1. Like many other E3 ligases, COP1 undergoes autoubiquitylation as a self-regulation mechanism (Bie and Ciechanover 2011; Seo et al. 2003). Here, UBP12/13 counteracted COP1’s self-degrading ability and positively promoted its activity. Our data support a mechanistic model where photoactivated CRY2 interacts with UBP12/13 DUBs, and this complex recruits COP1 by direct contact. We hypothesize that the UBP12/13 mediated deubiquitination of COP1 leads to its stabilization. The stabilized COP1 as part of the CUL4^COP1-SPA^ ubiquitin ligase then promotes ubiquitination and degradation of CRY2 in blue light (Figure 6). This dual mechanism of attenuation of CRY2 and stabilization of COP1 is unusual among the reported mechanisms but likely typifies mitigation of the receptor in an everchanging light environment of the plant. Such regulation by CRY2-UBP12/13-COP1 axis is particularly essential to keep a correct balance between the active and inactive CRY2 protein pool to optimize hypocotyl growth during photomorphogenesis.

**Figure 6.**
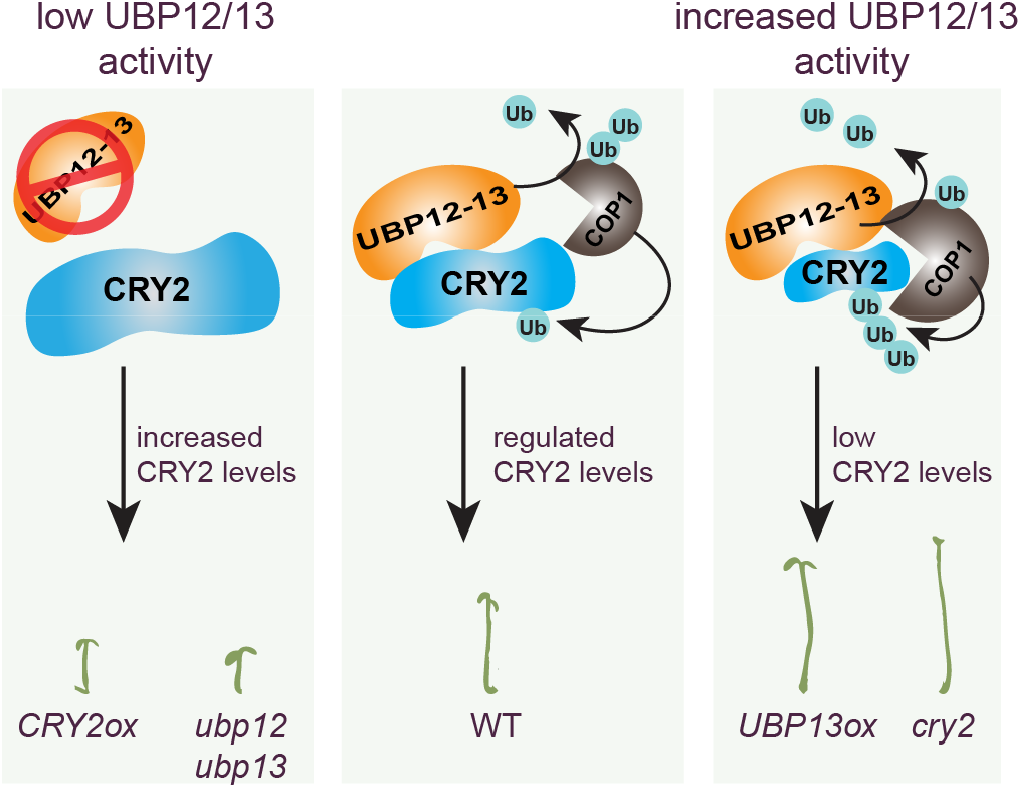
Model illustrating the role of UBP12 and UBP13 in regulating CRY2 protein and hypocotyl growth in blue light. During optimal photomorphogenesis (middle panel), CRY2-UBP12/13 complex recruits COP1 ubiquitin ligase and leads to its stabilization. This stabilized COP1 then targets CRY2 for ubiquitin-mediated proteasomal degradation leading to optimal hypocotyl growth. Low UBP12/13 activity seen in *ubp12ubp13* mutant, increase CRY2 levels due to lack of proteasomal degradation. Therefore, the hypocotyl phenotype of *ubp12ubp13* resembles *CRY2ox* (left panel). Accordingly, increased UBP12/13 activity results in enhanced ubiquitination and degradation of CRY2, as *UBP13ox* seedlings have a similar blue light insensitive phenotype as the *cry2* mutant (left panel).

Sufficient progress has been made on the specificity and regulation of deubiquitinase’s ability to identify different types of ubiquitin chain linkages and cleave them (Mevissen and Komander 2016). However, despite decades of research on deubiquitinases, how their catalytic activity and protein-substrate specificity are regulated is not well known. A handful of examples are available on the role of phosphorylation and ubiquitination in regulating the catalytic activity and localization of deubiquitinases. For example, phosphorylation of mammalian OTUB1, USP10, and ATXN3 results in their nuclear localization (Herhaus et al. 2015; Mueller et al. 2009; Yuan et al. 2010). But, how the activity of deubiquitinases is enhanced or inhibited by protein-protein interactions is poorly understood, especially in plants. Our results suggest that UBP12/13’s interaction with photoactive CRY2 regulates its activity and subsequent activation of COP1. Overexpression of COP1 in seedlings lacking UBP12 and UBP13 did not lead to the elongation of their hypocotyl compared to seedlings overexpressing COP1 alone (Figure 5E). Likewise, in the absence of UBP12/13, ubiquitination of CRY2 was absent (Figure 4D). This suggests that CRY2-UBP12/13 complex formation in light is required for the normal function of COP1, and UBP12/13 are themselves regulated by photoactivation of CRY2. Therefore, our results present an insight into how exogenous signals and their receptors regulate deubiquitinase activity through protein-protein interactions.

Cryptochromes, UBP/USP deubiquitinases, and COP1 are present in all major evolutionary lineages, pointing to a last common ancestor in which these proteins originated before plants and animals diverged (Liu et al. 2008b; Komander et al. 2009; Han et al. 2020; Cui et al. 2013). Importantly, light-dependent degradation of CRYs is preserved in flies, mammals, and plants to regulate their activity in timekeeping and growth (Peschel et al. 2009; Siepka et al. 2007; Busino et al. 2007; Godinho et al. 2007). However, the CRY2-COP1 interaction observed in plants is not conserved in mammals (Rizzini et al. 2019). In mammals, USP7 interacts with CRY1 and CRY2. But, contrary to our findings in plants, USP7 stabilized mammalian CRY1/CRY2 by deubiquitination (Papp et al. 2015; Hirano et al. 2016). Thus, the interaction between CRYs and the DUBS are mirrored in mammals and plants, but their roles and regulatory logic are reversed. Furthermore, contrary to the current assumption that deubiquitinases generally stabilize proteins, our unexpected findings accentuate the importance of deubiquitinases in controlling protein turnover and levels. Our study opens the door to future investigations on the deubiquitinase-dependent stabilization of ubiquitin ligases and their physiological roles in a multitude of developmental and growth programs.

## MATERIALS AND METHODS

### Plant material and growth conditions

Plants of Arabidopsis thaliana ecotype Columbia (Col-0) and mutant genotypes in this background were used. Arabidopsis mutants of *cry2-1* (Lin et al., 1998), *ubp12-1* (GABI_244E11), *ubp12-2w* (GABI_742C10), *ubp13-1* (SALK_128312), *ubp13-3* (SALK_130784), *ubp12-2w ubp13-3* (Cui et al., 2013), and *cop1-4* (McNellis et al., 1994) have been described previously. Transgenic line *cry2-1;UBQ10pro::9×Myc-6×His-3×Flag-CRY2* (*Flag-CRY2/Flash-CRY2*) has been reported previously (Pedmale et al., 2016). For all experiments, seeds were plated on 0.5× Linsmaier and Skoog (LS) medium (HiMedia Laboratories) with 0.8% agar, stratified for 2 days at 4°C in darkness and grown under the indicated light conditions in a LED growth chamber (Percival Scientific).

### Cloning and generation of Arabidopsis transgenic lines

All coding and promoter sequences were amplified from a cDNA pool from WT plants or a plasmid containing the CDS using standard PCR techniques and cloned in one of the Gateway donor vectors (pDONR-221, pDONR-P4P1R, or pDONRP2RP3) using BP Clonase II (Thermo Fisher). We used multisite Gateway cloning using LR Clonase II (Thermo Fisher) to combine the donor constructs with either pB7m34GW or pK7m34GW binary destination vector (Karimi et al., 2007) to generate the final expression constructs. *Agrobacterium tumefaciens* (GV3101) containing the expression constructs were used to transform Arabidopsis by floral dip method (Clough and Bent, 1998) to generate the transgenic lines. *Ubiquitin 10* (*UBQ10*) constitutive promoter was used to drive the expression of UBP13WT, UBP13-CD, and UBP13ΔMATH tagged with 6×HA epitope tag. UBP13-CD was generated by replacing Cys207 with a Ser residue in the UBP13-WT coding sequence by site-directed mutagenesis and UBP13ΔMATH line was generated by deleting the MATH domain residues 1-178 a.a. CRY2 overexpressing lines in *ubp12;ubp13* background was generated by transforming it with the *UBQ10pro::Flash-CRY2* construct. COP1 overexpressing lines in Col background was generated by transforming it with a *UBQ10pro::COP1-6xHis-3xFlag* construct generated as described above.

### Hypocotyl length measurements

The indicated genotypes were grown in 1 µmol m^-2^ s^-1^ constant blue light, 20 µmol m^-2^ s^-1^ constant red light or 100 µmol m^-2^ s^-1^ constant white light in a LED chamber at 22°C (Percival Scientific). Following the growth period, seedlings were placed horizontally and imaged using a flatbed scanner (Epson). Hypocotyl length was then measured using ImageJ software.

### Transient expression in *N. benthamiana* leaves

5 mL overnight cultures of *Agrobacterium tumefaciens* (GV3101) containing the expression constructs were centrifuged at 4000g at RT for 15 min. The bacterial pellet was then resuspended in infiltration buffer (10 mM MgCl_2_, 10 mM MES, pH 5.6, and 200 µM acetosyringone) and incubated at room temperature for at least 2 h. Each Agrobacterium suspension was then mixed in an equal ratio along with p19 (Lombardi et al., 2009) to a final OD600 of 1.0 before infiltrating the abaxial side of *N. benthamiana* leaves using a needleless 1 mL insulin syringe.

### Localization of UBP13-mCitrine and UBP13-ΔMATH-mCitrine

The UBP13-WT and UBP13ΔMATH were cloned into pDONR-221 vector. The *UBP13* promoter sequence used was the 2043 bp upstream of the start codon of *UBP13* ORF (Cui et al., 2013) and was cloned into the pDONR-P4P1R vector. These entry clones were then recombined with mCitrine (in pDONR-P2RP3) along with the pB7m34GW destination vector. The *UBP13pro::UBP13-mCitrine* and *UBP13pro::UBP13ΔMATH-mCitrine* constructs were then transformed into *A. tumefaciens* and infiltrated into *N. benthamiana* leaves. The plants were kept for 1-day in light and then 1-day in dark, and subsequently leaf discs were taken. The leaf discs were then counterstained by immersing them in 0.02 µg/mL DAPI in water for 3 min and rinsed 2× in water before imaging them using a fluorescence microscope (Evos, Thermo Fisher).

### Bimolecular fluorescence complementation (BiFC)

The *CRY2, UBP13* and *COP1* coding regions were cloned into pDONR-P2RP3 or pDONR221 vectors to obtain entry clones. These entry clones were then recombined together with either *UBQ10* or CaMV 35S promoter (in pDONR-P4P1R vector) and cVenus/nVenus (in pDONRP2R-P3 or pDONR-221) entry constructs along with pB7m34GW or pK7m34GW destination vectors using LR II clonase. The final expression constructs *35S::nVenus-CRY2, UBQ10::UBP13-cVenus, UBQ10::COP1-nVenus* and *UBQ10::nVenus-mock* were transformed into *A. tumefaciens*, mixed equally in the desired combinations and infiltrated into abaxial side of *N. benthamiana* leaves. Fluorescence signal was analyzed 2 days post infiltration (1-day light/1-day dark) on the abaxial side of leaf discs using a fluorescence microscope.

### Co-immunoprecipitation in *N. benthamiana* and *Arabidopsis*

For co-immunoprecipitation (co-IP) assays using *N. benthamiana*, the following constructs were transiently expressed in the indicated combinations by Agrobacterium infiltration and the leaves were collected after 3 days and flash-frozen using liquid nitrogen. The constructs used were *UBQ10pro::Flash-CRY2, UBQ10pro::Flash-mCitrine, UBQ10pro::UBP13-6xHA, UBQ10pro::UBP13ΔMATH-6xHA* and *UBQ10pro::COP1-6xHis-3xFlag*. For co-IP assays in Arabidopsis, stable transgenic lines expressing both *CRY2pro::Flash-CRY2* and *UBQ10pro::UBP13-HA* was used which was made by transforming *UBQ10pro::UBP13-HA* into *cry2;CRY2pro::Flash-CRY2* expressing line. 4-day old Arabidopsis etiolated seedlings were collected in dark or exposed to 30 µmol m^-2^ s^-1^ blue light for 10 min before sample collection.

Frozen plant tissue was first ground by mortar and pestle using liquid nitrogen and then resuspended in SII buffer (100 mM sodium phosphate, pH 8.0, 150 mM NaCl, 5 mM EDTA, 5 mM EGTA, 0.1% Triton X-100, 10 mM NaF, 1.5× protease inhibitor cocktail (Sigma), 20 µM bortezomib (MedChemExpress) and 20 mM N-ethylmaleimide). The extracts were sonicated at 30% amplitude, 0.5 S on/off for a total of 10 S and clarified by 2× high speed centrifugation for 10 min at 4°C. Proteins were then quantified using Bradford reagents with BSA as a standard. 2 mg of total protein was then incubated with anti-Flag antibody (M2 monoclonal clone, Sigma) for 1 h with gentle rotation at 4°C and then protein-G coated magnetic beads (Thermo Fisher) were added and incubated for an additional 30 min. The beads were then washed 3× with 0.8mL of SII buffer and the proteins were eluted from it using 2× Laemmli sample buffer by heating at 90°C for 5 minutes.

### Tandem Ubiquitin Binding Entities (TUBE) pulldown

Immunoprecipitation of ubiquitinated proteins from *cry2;UBQ10pro::Flash-CRY2* and *ubp12ubp13;UBQ10pro::Flash-CRY2* seedlings was performed as previously described with some minor modifications (Zhang et al., 2017). 4-day old etiolated seedlings were first pre-treated by transferring them to liquid 0.5× MS medium containing 0.01% Silwet L-77 and 20 µM MG132, then vacuum infiltrated for 10 min and kept in dark for 2 hours. Thereafter, the seedlings were either kept in dark or transferred to 30 µmol m^-2^ s^-1^ blue light for 30 min before being collected. Total proteins were using a buffer containing 100 mM MOPS, pH 7.6, 150 mM NaCl, 0.1% NP40, 1% Triton X-100, 0.1% SDS, 20 mM Iodoacetamide, 1 mM PMSF, 2µg/mL aprotinin, 40µM MG132, 5µM PR-619, 1 mM 1,10-Phenanthroline and 2× Complete protease inhibitor cocktail and PhosStop cocktail (Roche). 2 mg total protein was incubated with 30 µl agarose-TUBE2 or agarose-control beads (tebu-bio, Le Perray-en-Yvelines, France) for 5 h at 4°C with gentle rotation. The agarose beads were washed three times with the extraction buffer and eluted with 2× Laemmli sample buffer at 90°C. The eluate was used for immunoblot analysis using anti-Flag-HRP (Sigma) for detection of Flash-CRY2 and anti-P4D1-HRP (Santa Cruz Biotechnology) antibody was used to detect ubiquitinated protein. Anti-RPN6 antibody was used to monitor uniform loading.

### Immunoblot analysis

Proteins were separated in 10% Bis-Tris polyacrylamide gel using MOPS buffer and transferred to nitrocellulose membrane (Sigma) for immunoblot analysis. A polyclonal antibody against the C-terminal of CRY2 (SEGKNLEGIQDSSDQ) was produced in rabbit and used to detect CRY2 protein. The anti-CRY1 antibody (Liu et al., 2016) was obtained from Dr. Chentao Lin. anti-COP1 antibody to detect COP1 was described previously in Maier et al. (Maier et al., 2013). An anti-actin antibody (MP Biomedicals) was used as a loading control.

### RNA-sequencing (RNA-seq) and analysis

Seedlings were grown for 4 days in 1 µmol m^-2^ s^-1^ constant blue light and frozen in liquid nitrogen prior to RNA extraction. Total RNA was extracted using the RNeasy Plant Mini kit (Qiagen). 500 ng of total RNA was used to isolate poly(A)-mRNA using the NEBNext poly(A) mRNA Isolation Module (NEB) and the purified mRNA was used to build sequencing libraries using the Ultra II Directional RNA library Prep Kit for Illumina (NEB). Single end sequencing of 76 bp were performed on NextSeq instrument (Illumina). The sequencing reads were mapped to the *Arabidopsis thaliana* Col-0 reference genome (TAIR10) using the STAR aligner (Dobin et al., 2013). Differential gene expression analysis was performed using Cuffdiff version 2.1.1. for Linux (Trapnell et al., 2012). Gene Ontology (GO) term enrichment was performed on Panther Classification System (Thomas et al., 2003) using Fisher’s Exact test and Bonferroni correction. R environment (R Foundation) and its packages (ggplot2, ComplexHeatMap, RColorBrewer, gPlots, corrplot) was used for statistical analysis and to visualize results. De novo cis-motifs in the promoters were identified using HOMER (Heinz et al., 2010).

## ACKNOWLEDGEMENTS

We thank K. Schwartz for the technical support, plant care and propagation. We thank P. Sridevi, R. Martienssen, D. Jackson, Z. Lippman for critical reading of the manuscript. We thank C. Lin and U. Hoecker for the anti-CRY1 and anti-COP1 antibodies. O. Nilsson and Å. Strand for the scientific experimental support to L.N.L. This work is supported by National Institutes of Health (NIH) grant R35GM125003 to U.V.P. NIH GM12500303S1 and GM12500304S1 grant for the purchase of Zeiss LSM900 Airyscan2 confocal microscope is acknowledge by U.V.P.

## AUTHOR CONTRIBUTIONS

U.V.P. conceived the study. L.N.L and U.V.P. designed the experiments and L.N.L performed most of the experiments with the following exceptions: U.V.P. performed RNA-seq analysis, O.A. performed the CRY1 and COP1 protein analysis. A.A. performed phenotypic analysis. L.N.L. and U.V.P. wrote the manuscript, all authors reviewed and commented on the manuscript.

## Data and materials availability

Further information and requests for resources and reagents should be directed to and will be fulfilled by the Lead Contact, U.V.P.

## Supplementary Materials

Materials and Methods

Supplementary figures S1 – S5

Supplementary Table S1

