## Supplemental data for "UBP12 and UBP13 deubiquitinases destabilize the CRY2 blue-light receptor to regulate growth"

**The supplementary material includes:**

Materials and Methods
Figures S1 to S5
Table S1

### MATERIALS AND METHODS

**Phylogenetic analysis.** For phylogenetic analysis of the 27 UBP proteins in Arabidopsis, the full-length amino acid sequences were used. For phylogenetic analysis of UBP12/13 orthologs in other plant species, insect, vertebrate and worms, the amino acid sequence was obtained from NCBI. The alignment of the protein sequences was performed using ClustalW. The phylogenetic tree was constructed using MEGA-X version 10.1.8 by identifying conserved positions of the alignment using the Neighbor-Joining method with bootstrap test set to 1001 replicates.

**Affinity purification and mass-spectrometry.** 5-day-old *cry2-1;UBQ10pro::9×Myc-6×His-3×Flag-CRY2* (Flag-CRY2/Flash-CRY2) and WT control seedlings grown in white light were exposed to attenuated blue light conditions for ~16h as previously described (Pedmale et al., 2016). Total protein was extracted as previously described using SII buffer (Pedmale et al., 2016). Roughly 28mg of total protein was incubated with 150ml of protein-G-Dynabeads (Thermo Fisher) coupled to Flag antibody (M2 clone, Sigma). Protein complex was eluted with 3x-Flag peptide and precipitated with TCA. Liquid-chromatography coupled to mass spectrometry (LC-MS) was performed using standard techniques.

**Oligos used for genotyping and cloning.**

| Primer set | Forward | Reverse |
|---|---|---|
| *ubp12-2w*[1] | TGGTATGCCTTGCAGATTTTC | TTCATGTTTTGGGGCTAATTG |
| *ubp13-3*[2] | TTTACGATGCTGTCCTCGATG | AGCCATTTTATTCAATTGCCC |
| *cry2-1*[3] | GTTTCGTTAGTTCGGGACCA | CCGAAATGGAGATACGGAGA |
| UBP13 WT[4] | GGGGACAAGTTTGTACAAAAAAGCAGGCTccATGACTATGATGACTCCGCC | GGGGACCACTTTGTACAAGAAAGCTGGGTcATTGTATATTTTCACCGGCTTCTC |
| UBP13 ΔMATH[4] | GGGGACAAGTTTGTACAAAAAAGCAGGCTATGGCTGTGCGTAAAGTT | GGGGACCACTTTGTACAAGAAAGCTGGGTcATTGTATATTTTCACCGGCTTCTC |

| | | |
|---|---|---|
| COP1[5] | AAAAAGCAGGCTggATGGAAGAGATTTCGACGGATCCGG | AGAAAGCTGGGTgCGCAGCGAGTACCAGAACTTTG |
| UBP13 promoter[6] | GGGGACAACTTTGTATAGAAAAGTTGccTCCTCAAACCTCATTGAGGTTT | GGGGACTGCTTTTTTGTACAAACTTGgCATAGTCATTGCTCCGATAGTGAT |
| UBP12 promoter[6] | GGGGACAACTTTGTATAGAAAAGTTGggCACGATCTAAGTCAACAAATGTGAC | GGGGACTGCTTTTTTGTACAAACTTGgTGGCCGGAGAAGGATTAGA |
| attB1/B2 adapter primer[5] | GGGGACAAGTTTGTACAAAAAAGCAGGCT | GGGGACCACTTTGTACAAGAAAGCTGGGT |

1. Genotyping of GABI_742C10, WT 1244 bp, RP used with GABI-Kat LB for T-DNA insertion.
2. Genotyping of SALK_130784, WT 1118 bp, RP used with SALK LBb1.3 for T-DNA insertion.
3. Genotyping of *cry2-1*, WT amplifies ~550 bp and *cry2-1* mutant amplifies ~443bp.
4. Cloned with full length attB1 and attB2 sites.
5. Cloned with two step PCR attB1 and attB2 sites using the attB1/attB2 adapter primers.
6. Cloned with full length attB4 and attB1 sites.

### Lindbäck et al., Supplementary figure 1

**A**

| Accession | Protein name | Peptides (#) | Coverage (%) |
|---|---|---|---|
| AT1G04400 | CRY2 | 1222 | 89.70 |
| AT5G06600 | UBP12 | 80 | 39.6% |
| AT3G11910 | UBP13 | 57 | 29.4% |
| AT2G46340 | SPA1 | 56 | 29.4% |
| AT2G32950 | COP1 | 50 | 35.3% |
| AT4G08920 | CRY1 | 8 | 5.0% |
| AT3G52740 | BIC1 | 7 | 20.7% |
| AT1G53090 | SPA4 | 3 | 5.5% |
| AT3G15354 | SPA3 | 2 | 3.2% |

**B**

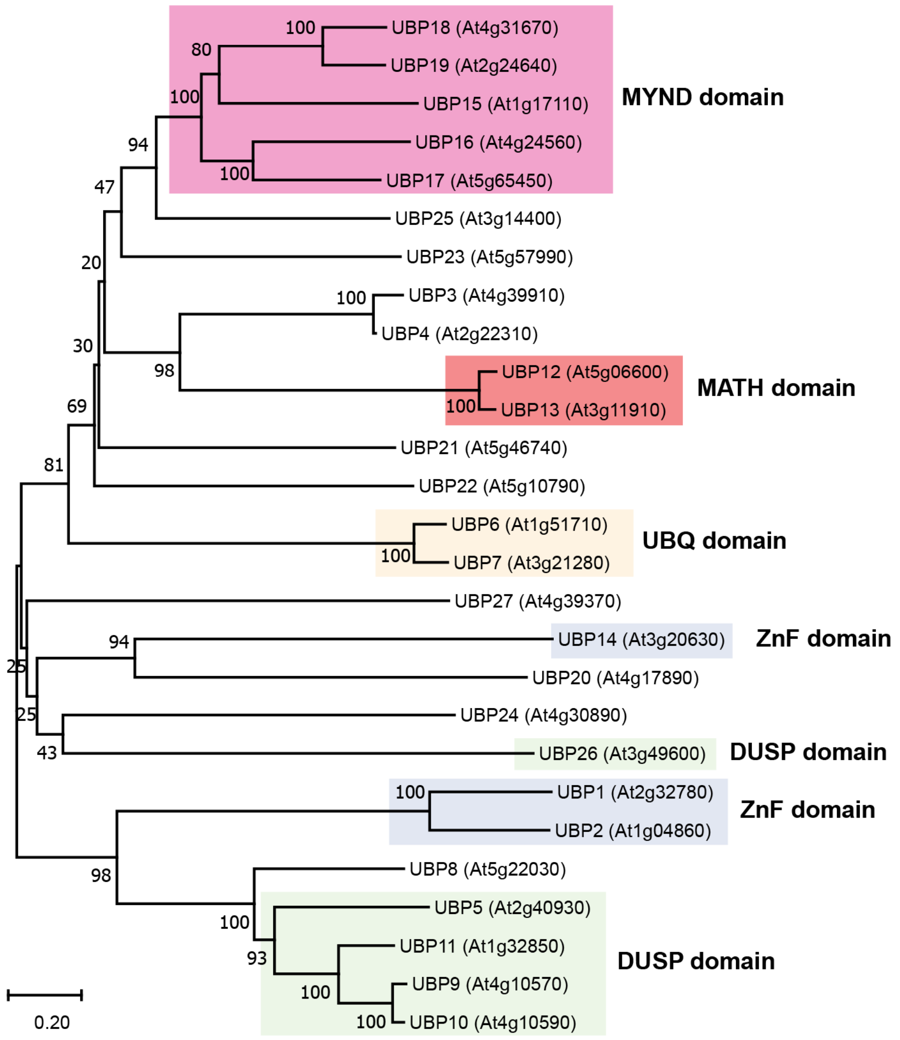

**Fig. S1.** (**A**) Summary of the *Arabidopsis* proteins including the peptide count and sequence coverage that form a complex with CRY2 as determined by affinity purification/mass-spectrometry (AP/MS) analysis under physiological conditions.
(**B**) Phylogenetic analysis of 27 UBP protein family members in *Arabidopsis*. The evolutionary history was inferred using the Neighbor-Joining method. The optimal tree with the sum of branch length = 18.12934367 is shown. The percentage of replicate trees in which the associated taxa clustered together in the bootstrap test (1001 replicates) are shown next to the branches. The tree is drawn to scale, with branch lengths in the same units as those of the evolutionary distances used to infer the phylogenetic tree. The evolutionary distances were computed using the Poisson correction method and are in the units of the number of amino acid substitutions per site. Abbreviations: UBP, ubiquitin specific protease; MYND, myeloid; MATH, meprin and TRAF homology; UBQ, ubiquitin homologues; ZnF, zinc finger; DUSP, domain in ubiquitin-specific proteases.

### Lindbäck et al., Supplementary figure 2

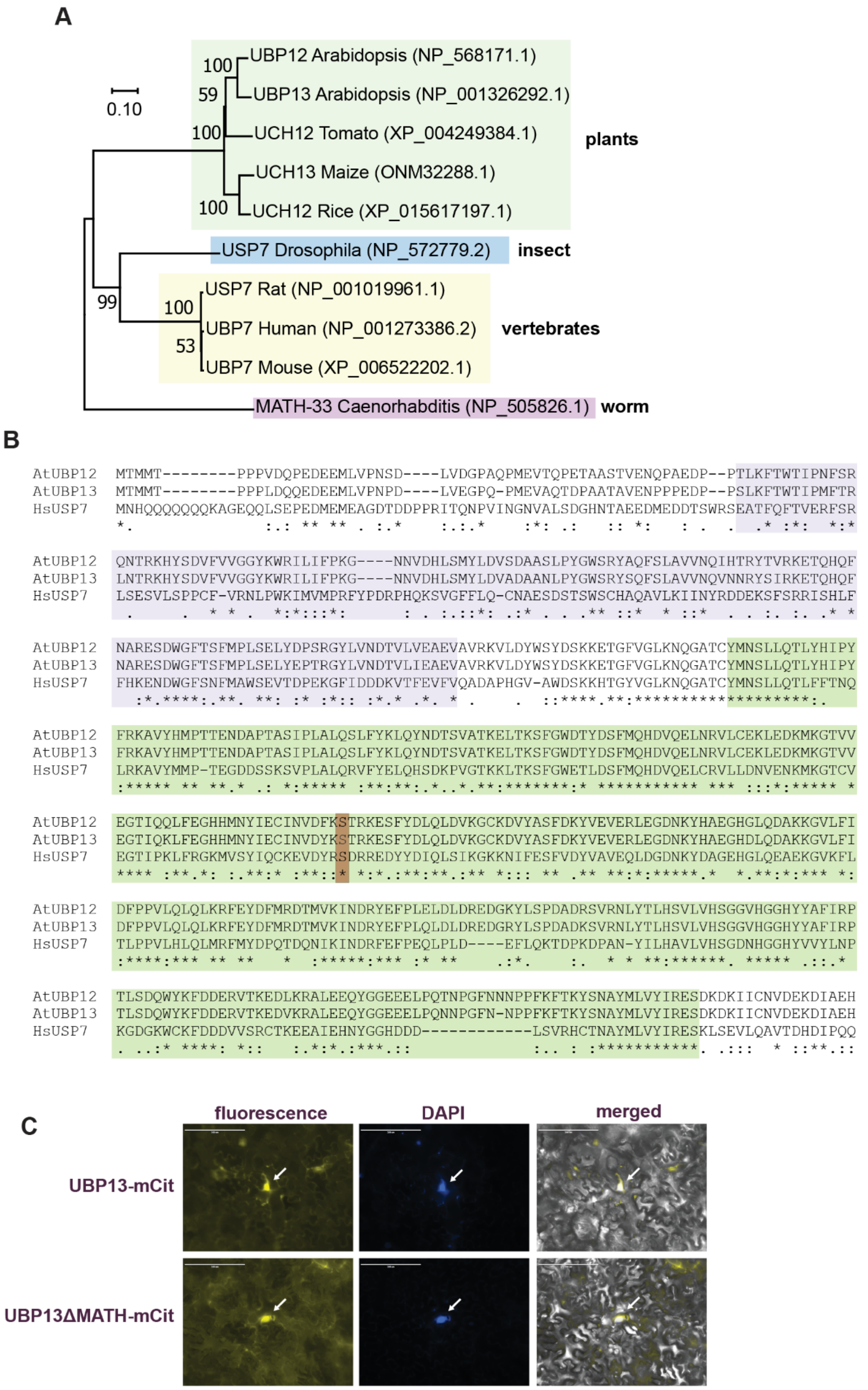

**Fig. S2.** (**A**) Phylogenetic analysis of *Arabidopsis* UBP12 and UBP13 protein along with their orthologs in major plant and animal species. The evolutionary history was inferred using the Neighbor-Joining method [1]. The optimal tree with the sum of branch length = 2.40603150 is shown. The percentage of replicate trees in which the associated taxa clustered together in the bootstrap test (1001 replicates) are shown next to the branches.
(**B**) Partial amino acid sequence alignment for the *Arabidopsis* UBP12, UBP13, and human USP7 deubiquitinases indicating the conserved domains and catalytic residues between them. Asterisks (*) indicate fully conserved amino acid residues, colons (:) indicate conservation between amino acid groups of similar properties, periods (.) indicate conservation between amino acid groups of weakly similar properties, and dashes (-) indicate gaps introduced to maximize alignment. The active cysteine residue is indicated by an orange box, the purple shade indicates the location of AtUBP13 MATH domain, and the green shade indicates the location of AtUBP13 UBP domain.
(**C**) UBP13 tagged with mCitrine fluorescent protein (*UBP13pro::UBP13-mCit*) or without its MATH domain (*UBP13pro::UBP13ΔMATH-mCit)* were transiently expressed in *Nicotiana benthamiana* leaves. Microscopic analysis shows their nuclear localization. DAPI was used as a counterstain to mark the nucleus. Arrows indicate fluorescence signal and DAPI stain in the nucleus.

**Lindbäck et al., Supplementary figure 3**

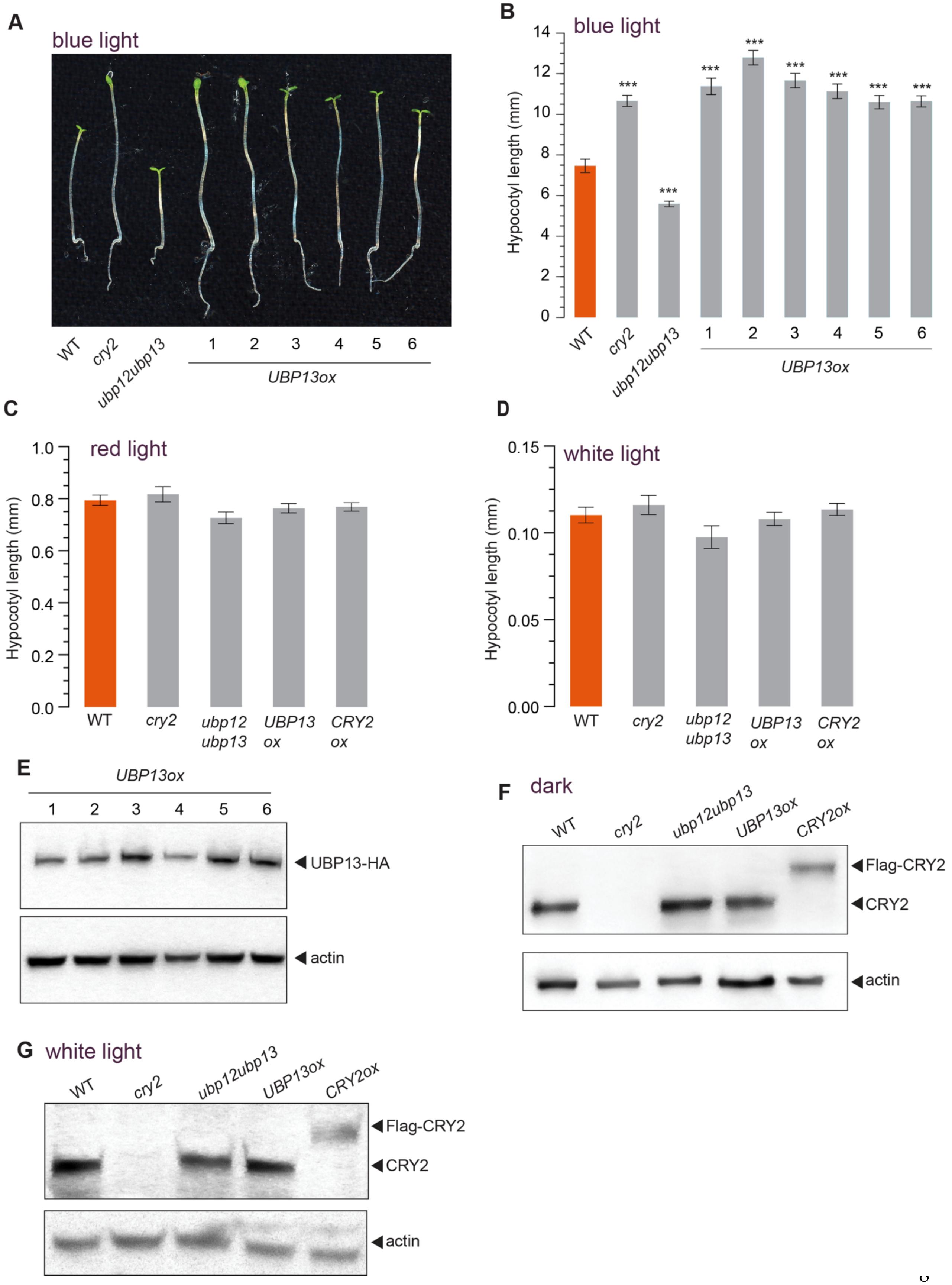

**Fig. S3. (A-B)** Hypocotyl length analysis of the indicated genotypes. Seedlings were grown for 4 days in 1 µmol $m^{-2}$ $s^{-1}$ constant blue light before measuring their hypocotyl length. Bars indicate average ± SE. Hypocotyl length was significantly different from wild type as demonstrated by Student's t-test: ***, $p < 0.001$.
**(C-D)** Hypocotyl length analysis of the indicated genotypes in red light (C) and in white light (D). Bars indicate average ± SE.
**(E)** Protein expression levels in independent transgenic lines expressing *UBQ10pro::UBP13-6xHA* (*UBP13ox*) after 4 days of growth in blue light (1 µmol $m^{-2}$ $s^{-1}$). The blot was probed using an anti-HA antibody. Actin is shown as a loading control.
**(F-G)** Seedlings of the indicated genotypes were grown for 4 days in dark (F) or 100 µmol $m^{-2}$ $s^{-1}$ white light (G) and analyzed for CRY2 protein levels using an anti-CRY2 antibody. Actin is shown as a loading control.

### Lindbäck et al., Supplementary figure 4

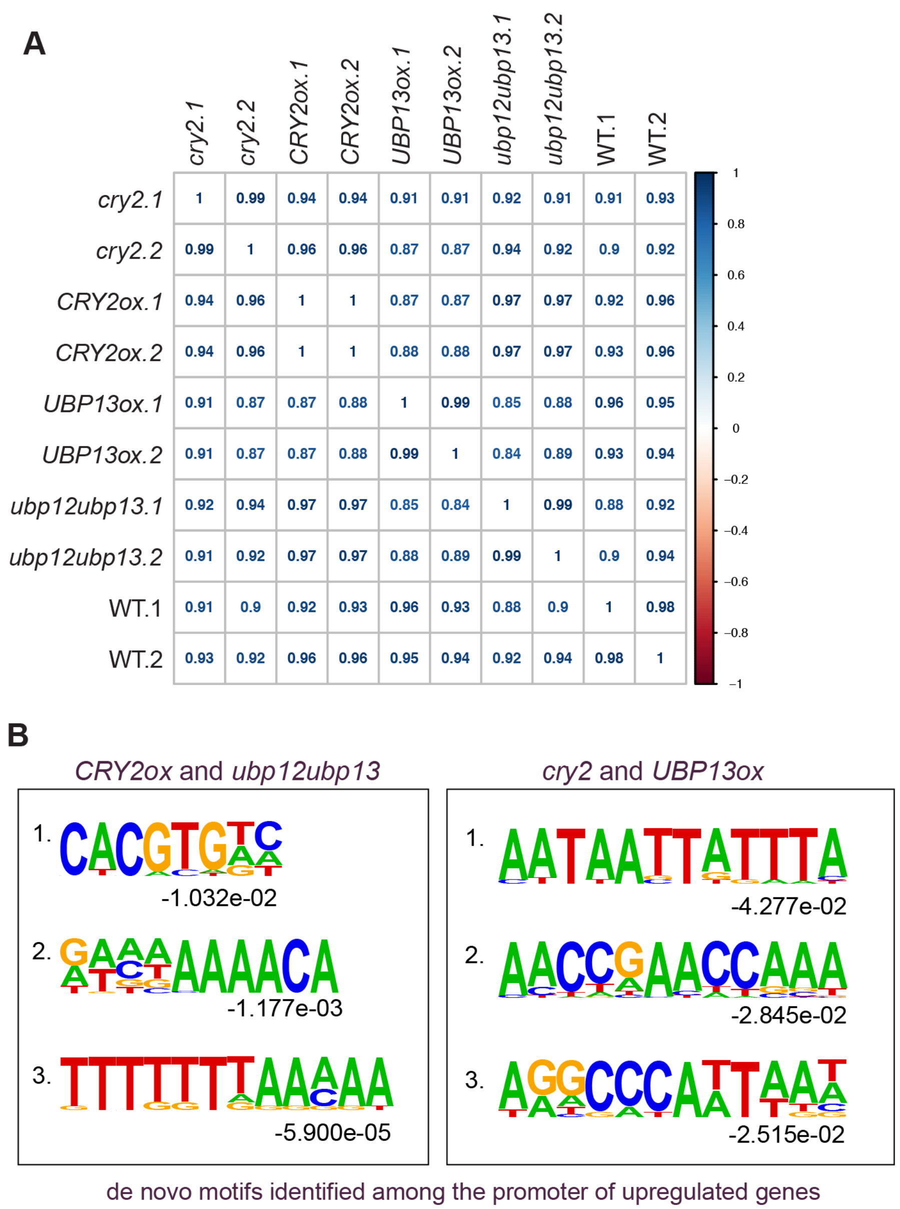


**Fig. S4.** (**A**) Pearson's correlation between the replicates used for the RNA-seq experiment. **(B)** Top 3 *de novo* motifs identified in the promoter sequence of common genes upregulated by the indicated genotypes. 500 bp upstream of the start codon of the gene was used to identify *de novo* sequences.

### Lindbäck et al., Supplementary figure 5

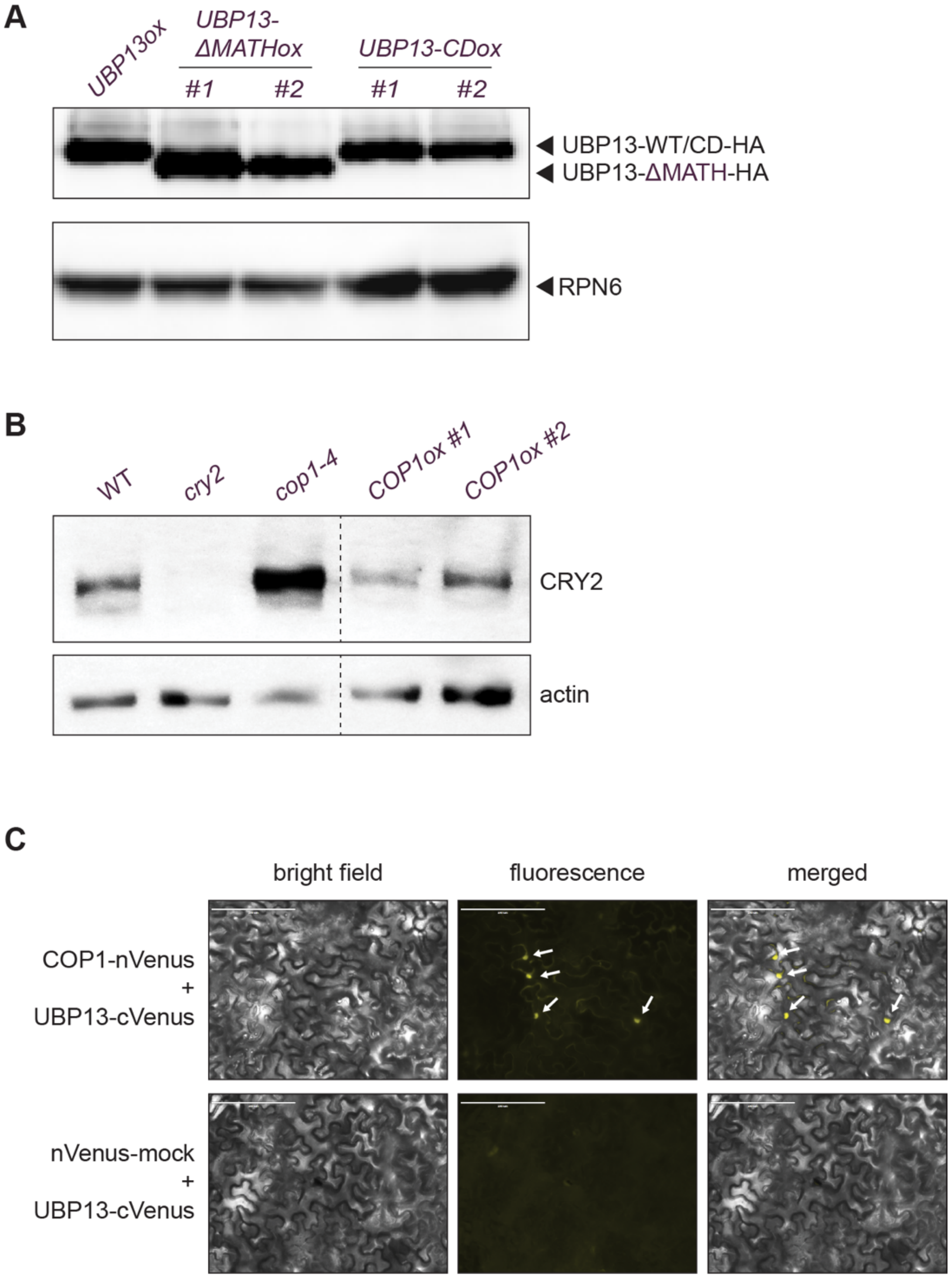


**Fig. S5.** (**A**) Protein expression levels in *UBQ10pro::UBP13-6xHA, UBQ10pro::UBP13ΔMATH-6xHA* and *UBQ10pro::UBP13CD-6xHA* expressing transgenic lines in *Arabidopsis*. The blot was probed using an anti-HA antibody. RPN6 is shown as a loading control.
(**B**) Seedlings of the indicated genotypes were grown for 4 days in 1 μmol $m^{-2}$ $s^{-1}$ blue light and analyzed for CRY2 protein levels using an anti-CRY2 antibody. Actin is shown as a loading control. The dashed line indicates that the blot has been cut and compared on the same blot.
(**C**) Interaction between COP1 and UBP13 was determined by bimolecular fluorescence complementation (BiFC) assay in *N. benthamiana* leaves. The arrows indicate fluorescence signal in a few selected nuclei of the epidermal cells.
